# Increasing protein stability by inferring substitution effects from high-throughput experiments

**DOI:** 10.1101/2022.05.18.492418

**Authors:** Rasmus Krogh Norrild, Kristoffer Enøe Johansson, Charlotte O’Shea, Jens Preben Morth, Kresten Lindorff-Larsen, Jakob Rahr Winther

## Abstract

Protein stability is an important parameter in almost all protein-engineering efforts. Evaluating the effects of the many possible amino acid changes to guide such projects is a significant task, even with recent advances in experimental and computational approaches. Here, we apply a computational model, GMMA, to extract substitution effects from a cost-effective genetic screen of a randomly mutated protein library. Using a high mutation frequency, the method can map stability effects of even very stable proteins for which conventional selection systems have reached their limit. Thus, we screened a mutant library of a highly stable and optimised model protein using an *in vivo* genetic sensor for folding and assigned a stability effect to 374 of 912 possible single amino acid substitutions. Combining the top 9 substitutions increased the thermodynamic stability by almost 50% in a single engineering step. This illustrates the capability of the method, which is applicable to any screen for protein function.

## Introduction

Protein engineering requires a complex concurrent optimisation of function, stability, and other desired traits, which can be cumbersome even with efficient screening automation. One reason for this is that many substitutions may be required to reach the desired phenotype, and the combinatorial space, when introducing multiple substitutions in protein sequences, quickly rises to, and above, experimentally accessible numbers. Efficient ways to navigate this space are therefore highly desirable^1^.

Even in the cases where enhanced stability is not the primary goal, it may still be a useful starting point when engineering enzymes^2^. As a general rule, proteins are only as stable as is required for the adequate fitness of their host^3^. It therefore stands to reason that it should be possible to stabilise most mesophilic proteins and enzymes for biotechnology purposes. This can provide the necessary stability head-room to alter a protein’s function, for example, where modification of a substrate cavity is suboptimal for stability^4^, or when multiple destabilising substitutions are required for directed evolution towards an altered function^5^. Thus, increasing the stability of a protein can increase tolerance to substitutions^6^, that might be needed for changes in the active site or other desired traits in protein engineering.

Directed evolution in combination with genetic selection and screening can address some of the challenges associated with the large sequence space. One of the tools that can be used to optimise stability are *Tripartite folding sensors*^7^. These are fusion proteins where the protein of interest (POI) is genetically inserted in a loop of a conditionally essential reporter enzyme^8^. Given a stable POI, the reporter enzyme is catalytically active, and the organism survives, while an unstable POI renders the fusion protein misfolded which abolishes the catalytic activity of the reporter thus impeding growth. Mutants can be selected for increased stability of the POI by increasing temperature^9^, antibiotic concentration^10^, or by following fluorescent readout^11^. Selection for aggregation-resistance^12^ and identification of stable protein scaffolds^13^ can also achieved using such systems. They will, however, eventually reach the limit of their dynamic range, requiring more complicated approaches to increase stability further^14^.

Genetic screening systems, when combined with massively parallel sequencing (MPS), can be exploited for protein science in range of powerful techniques broadly termed Deep Mutational Scanning (DMS)^15^. The potential for applying the method for engineering has been shown in the optimisation of a *de novo* designed influenza inhibitor^16^ and the identification of stabilising substitutions by studying epistatic effects between the binding capabilities of single and double mutants^17^. More quantitative analyses have been enabled by a thermodynamic model that considers doubly substituted protein variants to infer the effect of single amino acid substitutions on both protein-protein interaction and folding free energies^18,19^, which was later shown to match chemical unfolding stabilities well^20^. We have recently used a folding sensor based on a bacterial heat shock response^21^ to select for variants with improved thermodynamic stability of an already stable protein^22^. In line with this, we have shown that single substitution effects may be obtained by analysing the results of a DMS experiment with many diverse multiple-substituted protein variants, and suggested a global multi-mutant analysis (GMMA) for this task^23^. GMMA rests on the observation that enhancing amino acid substitutions, although they have no phenotype in a given assay alone, can be identified by their ability to compensate deleterious substitutions, which have a phenotype. Combining the information of phenotype (e.g. growth/no-growth) and genotype (mutations in the gene) in many multiply mutated variants, allows for assignment of effects of individual substitutions, even if they do not display a phenotype on their own. Specifically, the current implementation is aimed at identifying *generally enhancing* substitutions characterised by additivity when combined.

We have previously developed an *in vivo* tripartite folding sensor based on the enzyme *orotate phosphoribosyl transferase* (OPRTase), encoded by the *pyrE* gene in *E. coli*. OPRTase which is essential for pyrimidine biosynthesis^9^. Cells defective in this enzyme can only survive on minimal medium if a pyrimidine source, e.g. uracil, is added. We engineered a circularly permutated variant of OPRTase as a folding sensor, termed CPOP, where POI’s are inserted between the former N- and C-termini in the circular permutated enzyme. While the circular permutation is fairly unstable, it still complements a *pyrE* deletion, however, it becomes highly sensitive to the folding competence of the inserted POI. As a proof-of concept, we enhanced the stability of a marginally stable designed protein (called dF106^24^) through conventional directed evolution. The resulting protein variant, dF106-L11P-D83V, henceforth termed *enhanced dF106* (edF106), had a high stability of 48 kJ/mol. Thus, this variant could not be improved further in the CPOP system because it had already reached the upper limit of the dynamic range in the screen.

In the present work, we have further increased the stability of edF106 using CPOP in a DMS experiment on a library of more than 14,000 edF106 variants carrying on average 9 amino acid substitutions. GMMA estimates the additive effect of single amino acid substitutions on stability and function formulated as a fitness potential that relates to the assayed function^23^. Because CPOP reports the folding competence of a variant, we will in this work refer to the estimated fitness potential as stability. Thus, by linking a sequence to the growth/no growth phenotype, a stability effect was assigned to each of 374 substitutions. By introducing the nine top ranking substitutions from this single experiment, the stability was enhanced to almost 70 kJ/mol with minimal structural changes. These results demonstrate how GMMA is capable of accurately identifying stabilising substitutions for optimisation beyond the dynamic range of the assay.

## Results and discussion

### Resilience towards mutations reflects thermodynamic stability

We previously optimised the stability of the designed protein edF106 to the limit of the selective screen using the CPOP folding sensor^9^. To map the effect of multiple mutations and push stability of edF106 beyond the 48 kJ/mol, we generated variant libraries using semi-randomised oligonucleotides in the CPOP system (Figure 1a). If misfolded, this imposes uracil requirement on the cells and allows for genetic selection^9^ (Figure 1b). To generate a suitable dataset for application of the GMMA method, previous analysis had identified two key requirements for the variant library^23^: i) The library should encode a large number multiply-substituted protein variants, each carrying different combinations of amino acid substitutions all of which must be found in different contexts so that all are connected. In practice, this is achieved by having many fold more variants than unique substitutions. ii) Similar to chemical unfolding experiments, the inactivation transition should be well probed. This is achieved by, on average, having a number of substitutions per variant that is close to the number of substitutions required to inactivate the protein (Supplementary figure 1).

**Figure 1:**
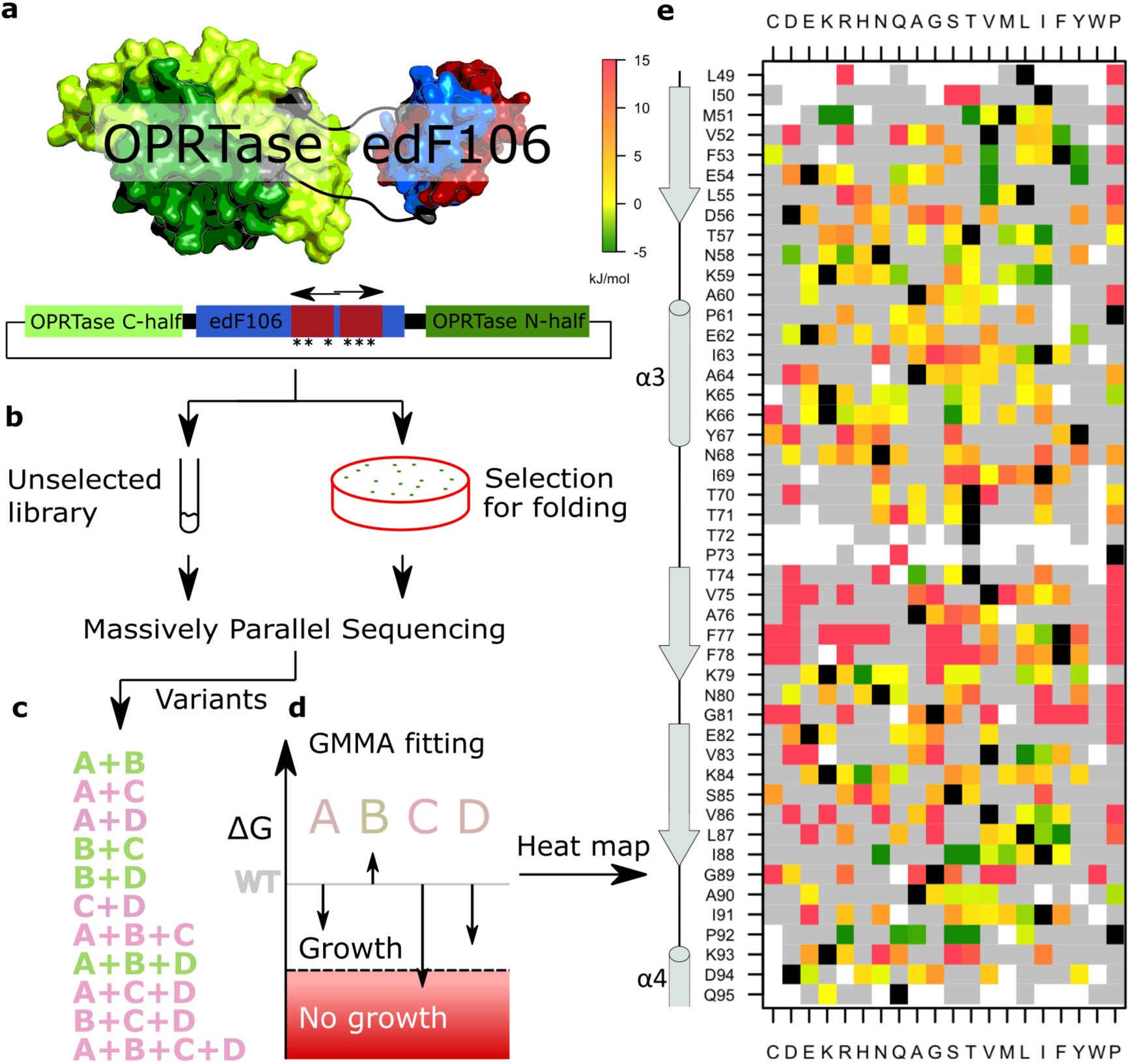
GMMA was used to analyse binary multi-mutant screening data from a genetic folding sensor and infer effect of individual amino acid substitutions. **a**, The enzymatic activity of the OPRTase (encoded by the pyrE gene) is essential for growth of E. coli on minimal medium and, in the CPOP system, this is dependent of the folding of edF106^9^. The C-terminal half (red) of the edF106 gene was mutated by PCR extension using long ‘doped’ primers (horizontal arrows). **b**, To screen variants for folding, the resulting plasmid libraries from unselected and selected cultures were subjected to MPS. **c**, The combined effect of multiple substitutions (here labelled A–D), complementing (green) and non-complementing (pink), was determined by comparing reads from the two libraries. **d**, The binary “growth/no-growth” in which variants are classified is modelled on a continuous and additive stability scale. Energies consistent with the growth data of the variants are indicated by vertical arrows. **e**, Heat map of the stability effects consistent with the combination of the substitutions in the library. Black squares represent “wild type” residues and grey squares represent substitutions observed in the libraries but in insufficient representation to obtain robust estimates, whereas white indicates that a variant is not found in the library.

Because the starting protein was already very stable (48 kJ/mol), we opted for a high mutation frequency obtained by using randomly mutated (‘doped’) oligonucleotides as degenerate primers with ∼10% error at each position of 77 and 75 bases length (Figure 1a). These were designed to cover the C-terminal half of edF106, amino acid residues 48-97, and to act as long primers for PCR amplification and assembled using USER cloning^25^. This approach was also chosen because previous work suggested that error-prone PCR may result in an overrepresentation of mutations that occur in early PCR rounds which is not ideal for GMMA^23^. Despite the huge diversity possible, we applied measures to obtain a library of limited size (about 10-20,000 variants) such that essentially all variants could be covered by MPS with a reasonable depth (see online methods). This was important because those variants present in the non-selected library, but not in the selected, were inferred to be non-functional in GMMA, and should therefore be identified with high accuracy (Figure 1b). MPS of the library revealed 15,018 unique DNA variants with, on average, 13.4 mutations per gene (∼9 amino acid substitutions), after processing the data and applying a quality cut-off on the reads (Supplementary Figure 2, Supplementary Table 1 and 2).

To identify the effect on stability of combinations of mutations using the CPOP system, the input library was plated on minimal medium to screen for growth (i.e. ura^+^ cells) and the resulting sub-library sequenced by MPS. From the input library, 19% of the sequences were recovered after selection at 30°C (Supplementary Figure 2). We emphasise that the edF106 protein is exceedingly stable and that the stringency of the screen was calibrated accordingly. This was also reflected in the survival of the variants, which depended on the number of substitutions in a slightly sigmoidal fashion instead of an exponential decay (Supplementary Figure 1). This is consistent with previous observations and indicated that the genetic data could be modelled by the thermodynamic stability effects of the protein^6^.

### GMMA analysis discovers stability effects

The principle behind the GMMA analysis is that the effect of a given amino acid substitution is determined in very many different variant contexts and thus that the inferred effect is mostly independent of a specific context. Such generally-stabilising substitutions may be recognised by their ability to compensate other destabilising substitutions. To illustrate this idea, consider two mildly destabilising substitutions, A and D, a stabilising substitution, B, and a deleterious destabilising substitution C (Figure 1c and 1d). Singly-substituted variants of A and D both show wild-type like growth (within noise) whereas they may both be inferred to be destabilising from the observation that the double-substituted variant A+D does not complement growth. On the other hand, a variant, A+B+D, with an additional substitution, B, is observed to complement growth from which we may infer that B is stabilising because it rescues the inactivation by A and D. Using this concept, the effect of each individual substitution is determined from combinations with many others in a global fit to all variants.

GMMA was essentially computed as described previously^23^. We used binary classification of folding and mis-folding and relied on the robustness of GMMA to infer a quantitative stability effect. We validated this approach by comparing our previous GMMA analysis to one in which we had artificially made the data binary and find that the two are strongly correlated (Pearson correlation of 0.97; Supplementary Figure 3. By using a binary phenotype, GMMA simplifies to a logistic regression model with the particular fitting scheme described previously. Briefly, initial estimates were obtained using the mean-field approach followed by a global optimisation using Levenberg-Marquardt damped least-squares. Reliable stability effects could be assigned to 374 out of the 838 unique substitutions in the library (Figure 1e).

### Stability measurements validate GMMA

To gauge the accuracy of our GMMA of the data from the folding sensor, we examined how well it could pinpoint the, presumably, rare stabilizing substitutions in edF106. As substitutions in proteins are on average mostly destabilizing^26–28^, this was a stringent test of our analysis. We therefore introduced the 15 substitutions predicted to be most stabilizing by the GMMA into the isolated edF106 protein to measure their individual as well as combined effects on thermodynamic stability (Figure 2a). We used a “two-dimensional” unfolding approach, where denaturant unfolding is combined with a temperature scan^29^, to measure the stability. For very stable proteins this analysis arguably gives a more accurate measure of stability because the unfolding transition is probed at different temperatures increasing the confidence in the extrapolation to a solution with no denaturant (Supplementary Figure 4 and 5). The top 15 substitutions are found at nine distinct positions, with some positions having two or three substitutions that GMMA suggests to be stabilizing. Thus, only nine could be combined to form the multi-mutant 9 (MM9) variant. To evaluate additivity, we constructed two additional multi-mutants with three and six of the nine top substitutions (MM3 and MM6). Interestingly, the multi-mutants show progressively increasing stability, whereas the individual stability effects vary more (Figure 2a). We compared the measured stability of MM3, MM6 and MM9 with the stability expected if all variant effects were additive and find that the total stability gains were 69% of the values expected (88%, 74% and 69% in MM3, MM6 and MM9, respectively.) All the same, the MM9 variant had a stability of 68.6±1.1 kJ/mol (ΔΔG =21.8±1.1 kJ/mol, n = 5) and an extrapolated melting temperature of 152±10°C (n = 5) (Figure 2b and 2c). While these values are somewhat uncertain due to the very long extrapolations to denaturant-free conditions, it is remarkable that this increase in stability was achieved in a single experiment of screening with a protein which was already very stable and which had hit the ceiling of the selection system used.

**Figure 2:**
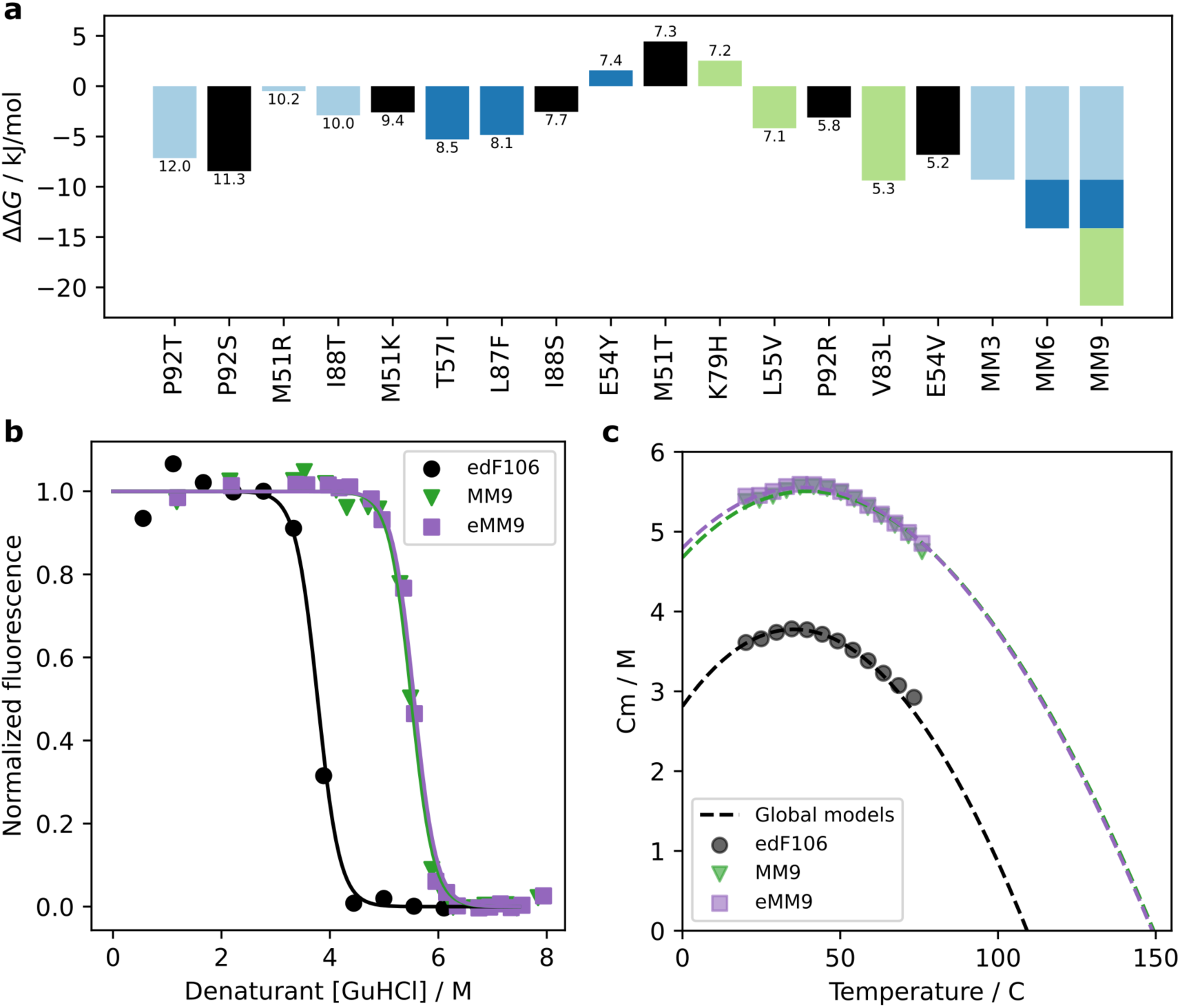
GMMA analysis successfully identifies substitutions that increase stability. **a**, Estimates of the change in stability for all single substitutions and combined multi-mutants (n = 1). The bars in the plot are coloured in light blue, darker blue and green to indicate their relationship with the MM3, MM6 and MM9 multi-mutants. Substitutions shown with black bars were not used in these multi-mutants because the GMMA analysis predicted that a more stabilising substitution was available at the same position. The magnitude of the contribution of each substitution in the stacked bars plotted for MM6 and MM9 is only to guide the eye and contains no information of the individual contributions from each substitution. Numbers below/above the bars indicate the stability predicted by GMMA. **b**, The average normalised fluorescence value of folding is shown as a function of denaturant for the starting point edF106 (black) and the stabilised multi-mutants: MM9 (green) and eMM9 (blue). **c**, The temperature dependence of the midpoint in the chemical denaturation, Cm, of the three proteins obtained by regular chemical denaturation fitting using isothermal slices of the NanoDSF data is plotted with the dashed line representing the global fit. The extrapolation of these fits to the absence of denaturant yields a melting point (temperature at which Cm=0) increase to ∼150°C. Illustrations in **b** and **c** is from the MM9_3 and eMM9_3 replicate and fits for the other replicates can be found in supplementary figure 4.

While 12 of the top 15 variants identified by GMMA were stabilizing in thermodynamic measurements of the isolated edF106 protein, three substitutions did not increase stability. Here, we must take into consideration that mutations were scored in the CPOP system where insertion of target protein into a fusion with the sensor protein is not completely equivalent to that of the target, in this case a thioredoxin domain, on its own. Notably, the poorly predicted positions, E54Y, K79H and M51T, are close to the N- or C-termini in the crystal structure of dF106 and are therefore in close physical proximity to the CPOP fusion site. Thus, mutations which could be stabilizing in the fusion might behave differently outside of the CPOP context. As two substitutions (E54Y and K79H) appeared to destabilise edF106 (Figure 2a, Supplementary Figure 4, Supplementary Table 3) and the optimal substitution appeared not to have been chosen at all positions, we prepared an enhanced version of MM9, termed eMM9, in which the best possible substitutions at each position, according to the measurements of single variants, were chosen: M51K, E54V, L55V, T57I, V83L, L87F, I88T, P92S. Within experimental error, however, we were not able to differentiate the stability of eMM9 from MM9 (Figure 2b and 2c) (ΔG = 69.5±1.1, n = 4, *p-value* = 0.30 (two-sided independent t-test)), again possibly due to the very long extrapolations to denaturant-free conditions. All things considered, effects on stability in the CPOP fusion are surprisingly well reflected in the isolated edF106 protein, and we note that including slightly destabilizing substitutions, as measured in single substitutions in edF106, did not abrogate the stabilization in the multi mutant context.

### Crystal structures show increased similarity to the 1FB0 design template

To gain more insight into the effects of the mutations, we crystalized and determined the crystal structure of MM9 using diffraction data that extended to a resolution at 1.9Å (Figure 3). Super imposition with the originally redesigned protein dF106 (PDB: 5J7D) showed a good fit with a RMSD of 1.20 Å using the C_α_ positions. However, there was no visible electron density to fit the N-proximal residues (1–17). Instead, a strong crystal contact was present on the exposed surface and had likely moved the 17-residue helix (α1) out of the way (Supplementary Figure 6). The crystals grew very slowly, typically within a month, with a low success rate out of 48 examined crystallisation conditions. Only one or two would yield crystals indicative of a conformational change taking place in order to stabilise the crystal lattice. The similar construct of eMM9, however, readily formed crystals in several conditions, with the best dataset collected to 2.0 Å resolution. The complete model could readily be built into the electron density despite minor crystal twinning present in the dataset that challenged the data analysis (Methods). This revealed an equivalent structure to MM9 with all C_α_ RMSD at 0.62 Å, whereas when structural compared to dF106 the RMSD was measured to 1.92 Å. With the N-proximal helix visible (α1), the main difference between eMM9 and the original designed protein, dF106, is indeed found in the in this region where eMM9 showed much better agreement with its original design template spinach thioredoxin (PDB: 1FB0). Interestingly, the other main region of change is the start of helix 2 (α2) – the active site of the spinach thioredoxin – where MM9/eMM9 is more akin to the natural protein. Thus, an all-C_α_ comparison of eMM9 showed a RMSD of 0.97 Å to spinach thioredoxin instead of 1.92 Å when compared to dF106 (Figure 3a). The structure also revealed that the destabilising substitution E54Y presented in MM9 but not in eMM9 would probably be incompatible with the native conformation of Val2 which may have facilitated its increased dynamics following MM9 crystal formation (Figure 3c). That this clash happens close to the N-terminal further strengthens the idea that the discrepancy is likely an artefact from the fusion protein and not the GMMA method.

**Figure 3:**
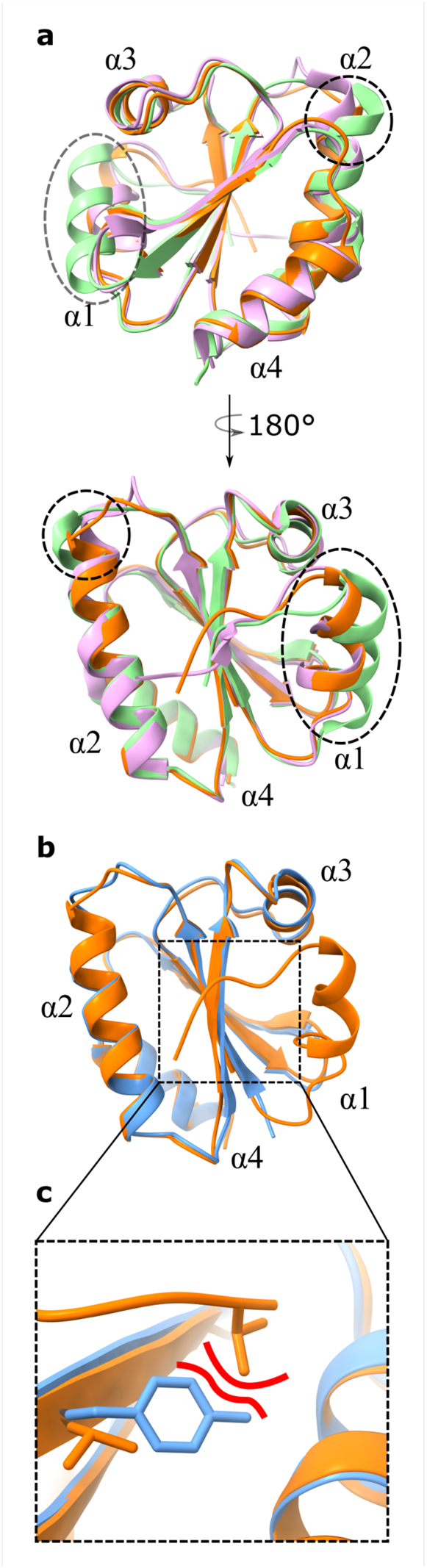
Crystal structure of the multi mutants shows increased template similarity. **a**, Cartoon rendering showing dF106 (PDB: 5J7D) in green, eMM9 (PDB: 7Q3K) in orange and the original spinach Trx design template (PDB: 1FB0) in purple. Bottom panel shows the structure flipped 180° around the vertical axis. Regions of structural re-arrangement are highlighted with in black circles. The mutated residues can be found drawn on the models in Supplementary Figure 6. **b**, Alignment of the structures of MM9 (PDB: 7Q3J, blue) and eMM9 (PDB: 7Q3K, orange) with **c**, showing in a magnified view that the placement of Val2 is incompatible with tyrosine at position 54.

### Structure and sequence-based methods do not predict most stabilizing variants

In general, we found it difficult to rationalise why many of the substitutions stabilised the protein. We therefore asked whether it would have been possible to point them out as likely candidates beforehand. We calculated variant effects using two methods; structure based stabilities calculations using Rosetta and a direct coupling analysis (lbsDCA) of a multiple sequence alignment (MSA) of natural thioredoxins as such analyses have previously been shown to predict stability effects^22,30,31^ (Figure 4). While both methods gave rise to reasonable overall Pearson correlations of 0.54 and 0.59 respectively, they failed to identify many of the substitutions which were inferred to be stabilizing (negative ΔΔG-values) in our GMMA. Some of the best substitutions found here score poorly in Rosetta, e.g. P92T and P92S have rank >50 and V83L, L87F and L55V rank >150, and all five scored to be destabilizing. The MSA based method, lbsDCA, in general performs better but still have V83L, P92S and P92T with rank >30 and E54V and L87F with rank >100.

**Figure 4:**
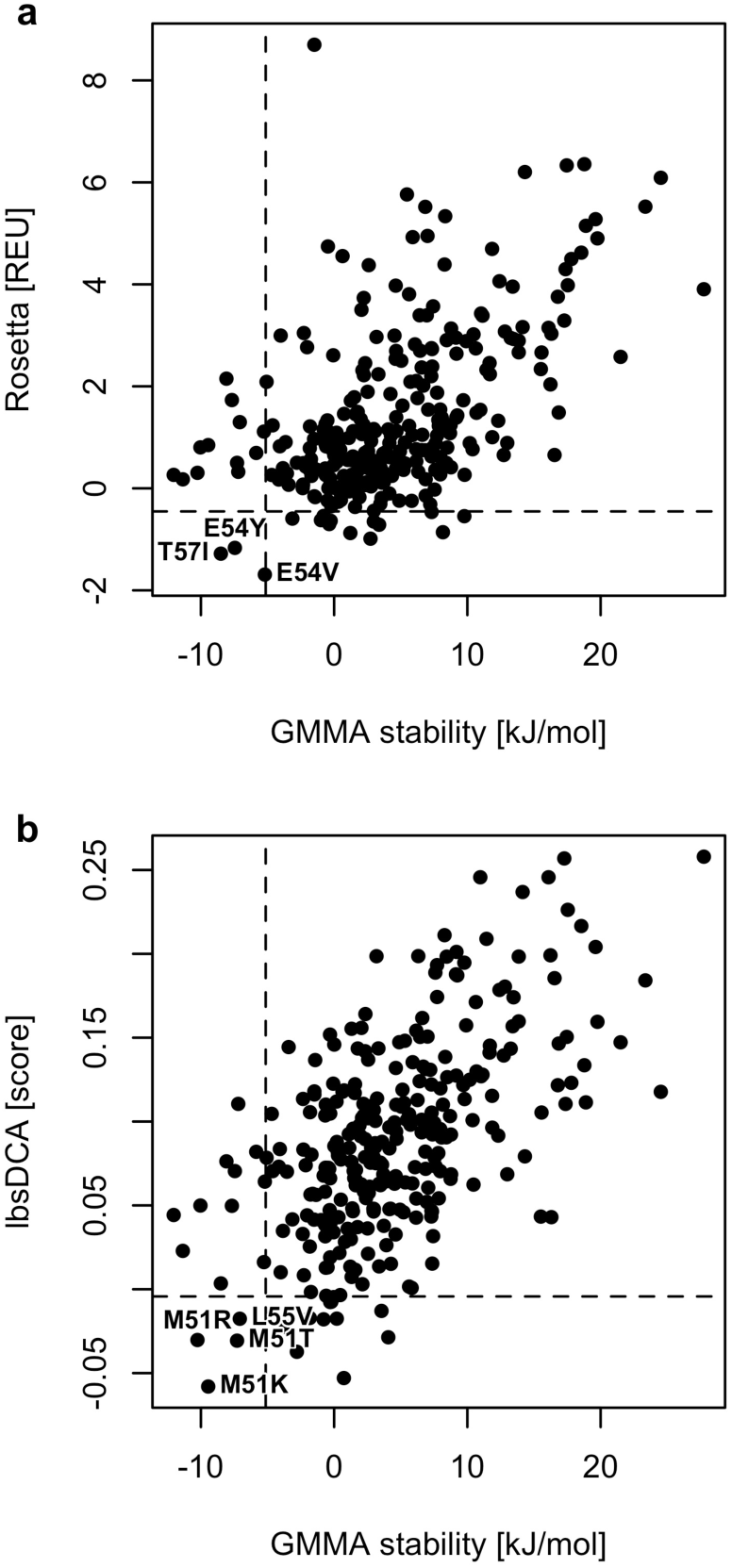
In-silico models inadequately recapitulates the GMMA-inferred stabilities. The correlation between the GMMA and stability predictions using Rosetta molecular modelling **a**, and an evolutionary conservation score, lbsDCA, in **b**, are shown (see methods). Pearson correlations are 0.54 and 0.59 in panels **a** and **b** respectively. The dashed lines indicate the value of rank 15 for each measure and show the overlap in the top 15. In general, the in-silico models do not reproduce the GMMA top15 well, highlighting the benefit of genetic screening coupled with GMMA in stability design efforts.

With 12 out of 15 substitutions confirmed to be stabilizing (figure 2) and the zero-effect accurately reproduced (Supplementary Figure 3), GMMA seems to identify stabilizing substitutions accurately. On the other hand, only 8 and 11 of the 15 top ranking substitutions from Rosetta and lbsDCA are estimated to be stabilizing respectively, and these effects are significantly smaller in magnitude as evaluated by GMMA. In particular, they identify only 3 and 4 of the GMMA top 15 ranking substitutions, respectively, and Rosetta rank 6 (K66I) and 13 (T74I) are estimated to be highly destabilizing and in the worst quartile of GMMA stabilities. This indicates that screening of multi-mutant coupled with GMMA is indeed a complementary and relevant method to obtain stability estimates for protein engineering.

## Conclusions

The GMMA approach shows remarkable efficiency in identifying the rare stabilising substitutions of the model protein edF106, allowing us to improve the stability by close to 50% in a single iteration. It is worth noting that the starting protein, edF106, can sustain growth even at the highest screening temperature 42°C, in the CPOP system, making it impossible to select for stability-enhancing substitutions simply by directed evolution^9^. Nevertheless, the selection for folding was here done under “mild” conditions (30°C), where the lower temperature imposes a less stringent conditions for protein folding. This highlights that the GMMA approach allows us to identify stability-enhancing substitutions outside the dynamic range of the selection. While other measures can be taken to address this issue^14^ we believe that the GMMA approach may be more robust. As GMMA only finds substitutions that are stabilising in many variant backgrounds, stabilising substitutions can be combined, suggesting that GMMA can efficiently guide optimisation in the vast combinatorial space of the protein sequence. For the same reason, we expect that interrogating parts of target proteins and combining information from fragments provides a useful and experimentally less demanding access to protein engineering. While the CPOP system has proven very successful here as a proxy for stability of a protein with no intrinsic function, GMMA can be applied to any system in which functional selection can be carried out in high throughput.

## Acknowledgements

We would like to acknowledge the help from Joseph Nesme and Shashank Gupta with obtaining the MPS data. We also thank the MAX-IV synchrotron staff, Ana Gonzales, for help during the remote X-ray diffraction data collection. We thank Matteo Cagiada and Anders Frederiksen for implementing computational pipelines for conservation and stability scores respectively. We are thankful for discussions with Associate Professor Martin Willemoës, and we are especially grateful for the help and funding provided by Professor Alexander K. Buell (Grant number: NNFSA170028392) to finalise the project.

## Data and code availability

All data and software generated for this study are available in the GitHub repository, https://github.com/KULL-Centre/papers/tree/main/2021/trx-gmma-norrild-et-al.

## Online methods

### Library construction

Libraries were constructed by using long “doped” oligonucleotides as primers for inverse PCR on a plasmid^9^ containing the fusion between the gene encoding CPOP sensor and a gene encoding edF106. Primers, obtained from LGC Biosearch technologies, contained a deoxyuracil at position 6 from the 5’ end so as to allow for annealing using a USER cloning approach^32^. Primers were named based on the amino acid positions mutated; oligo [48:72] and oligo [74:97], and all positions not involved in the USER-cloning site, 71 and 69 bases respectively, were “doped” with 10% of the three non-wild-type nucleobases for random mutagenesis. A version of the plasmid pMMA010^9^ with edF106 inserted in the CPOP system was amplified in a PCR reaction using PfuX7 polymerase, which is compatible with the USER-cloning^33^. 50 µL of PCR reaction mix [1x HF buffer (Thermo Fischer), 0.2 µM of each primer, 50 µM dNTPs, 0.15 ng/µL template plasmid] were prepared with 1 µL of in-house purified PfuX7. The reaction was run with initial denaturation at 98°C for 30 seconds and 30 cycles of 10 seconds at 98°C, 30 seconds at 62°C and 5 minutes at 72°C. A final 10 minutes at 72°C was employed to complete any unfinished product. Twenty 50 µL PCR reactions were pooled and the intended product was purified by gel band excision from a 1% agarose gel. DNA was extracted from the gel with a gel band purification kit (GE healthcare) and eluted from the spin columns of the kit with 50 µL MilliQ water. 100 µL reactions were prepared for USER excision and ligation consisting of 85 µL purified PCR product, 10 µL 10X T4 DNA Ligase buffer (Thermo) and 5 µL USER enzyme mix (New England biolabs)^34^. The temperatures used for the reaction were 1 hour at 37°C for catalysis, 30 minutes at 25°C for dissociation of excised fragment and annealing consisting of 20 minutes at each of the following temperatures: 12°C, 11°C, 10°C, 9°C and 8°C. The solution was kept on ice before immediately adding 5 µL of T4 DNA ligase (Thermo) and 10 µL 5 mM ATP (VWR Life science). The solution was then incubated for 30 minutes at room temperature before heat inactivation of the ligase at 70 °C for 10 min as recommended by the supplier. 2.5 µL Dpn1 (Thermo) were added before incubating the solution at 37°C overnight. Next day, the DNA was purified using the Illustra GFX PCR DNA and Gel Band Purification Kit (GE healthcare) and eluted in 20 µL sterilised MilliQ water.

### Initial library transformation

10 µL of purified cloned DNA was used to transform electrocompetent MC1061 cells ^35^ and the transformants were plated on four large 140 mm petri dishes to maximize colony separation, yielding an estimated 99,000 colonies. Colonies were scraped off the plates and collected in 5 mL sterile PBS buffer each (20 mM phosphate and 150 mM NaCl, pH = 7.4). The density of the recovered cells where normalised to OD_600_=3 before isolating plasmids using a mini prep kit (Omega) for each plate and eluting in 50 µL TE buffer (10 mM Tris-HCL and 1 mM EDTA, pH=7.3). To normalise the number of variants obtained from each purification, the purified plasmids were mixed in equimolar volume based on their absorbance at 260 nm.

### Retransformation

The plasmid libraries were transformed into the selection strain by using 5 ng of the purified and mixed plasmid library to transform electrocompetent MRH205 cells^9^. Ten-fold dilution of the transformed cells were plated on a 140 mm LB agar plate with 100 µg/mL ampicillin and 50 µg/mL kanamycin for a limited library size of estimated 53,600 colonies. Cells were collected using the same protocol as for the initial library transformation. A freeze stock was prepared for each library with 800 µL cell suspension and 200 µL of 87% glycerol. 50 µL aliquots were made from the freeze stock for screening of the libraries. Plasmids from single colonies of the library were purified and Sanger sequenced to confirm correct assembly of the library. Sequences appeared clear and uniform, suggesting that none of the transformants tested carried significant levels of more than one variant.

### Screening of library

The cell library was diluted 100,000-fold in PBS buffer and plated on three 140 mm selective medium plates at 30°C ensuring no more than 80 CFU/cm^2^ but a five-fold sampling depth of the library. After 22.5 hours incubation, plasmids from the plates were purified similarly to the collection of the initially transformed libraries. The concentration of DNA in the mini preps were normalised by dilution in buffer before mixing equal volumes of the solutions. In parallel, the library was also grown over night in 5 mL LB with 100 µg/mL ampicillin and 50 µg/mL kanamycin before purifying the plasmids of the full library using the same mini prep kit.

### Massively parallel sequencing

Sequencing of the purified plasmids were done with paired end amplicon sequencing protocol^36^ on one third of a Illumina MiSeq run using the version 3 kit with 600 cycles. The mutated part of the protein and 70 base pairs flanking region in each direction was sequenced. Amplicons with Illumina Nextera primers (361 base pairs) were produced by a PCR reaction with two HPLC purified primers with one part complementary to the plasmid and the Nextera sequence as a 5’ overhang. 25 ng template were used for the 25 µL PCR mix using HiFi Pfu polymerase (PCR biosystems). The reaction (Initial denaturation: 1 min at 95°C. Cycle: 15s at 95°C, 15s at 55°C and 1min at 72°C) was run for a minimum amount of cycles (12 cycles) as to reduce PCR chimeras^37^. The final elongation step was omitted to reduce the amount of chimeras^38^. Amplicons from the first PCR were purified from 20 µL using AMPure XP magnetic beads (Beckman Coulter) and eluting in 40 µL elution buffer. 2 µL of purified amplicons were used as templates for the second PCR to attach indexing primers to the Nextera adapters (total of 429 base pairs), including one reaction without template as a negative control. The PCR reaction was run for 15 cycles and was checked for uniform bands on a gel. 30 µL were purified using the magnetic beads (AMPure XP, Beckman Coulter) and eluted in 40 µL elution buffer. The concentrations of the samples were then normalised with SequalPrep Normalization Plate (Invitrogen) and eluted in 20 µL elution buffer. The amplicons were pooled and then cleaned and concentrated with the DNA Clean & Concentrator kit (Zymo). The concentration of the DNA in the eluate was quantified with Qubit and the sample was then diluted to 4.5 nM. The sample was denatured and diluted as described by the Illumina protocol for the MiSeq system. 200 µL sample were mixed with 400 µL of other samples for the run before removing 30 µL of the pooled samples and adding 30 µL PhiX (5% spike) as internal control. 1.35-1.5 million reads were collected totalling ∼3 million paired end reads before and after selection (Supplementary Table 1).

### Processing the paired end reads

Data from the sequencing were demultiplexed and trimmed to the start of the plasmid coding sequence. Filtering was done with a custom python script (`Filtering.py`) that checked for full complementation of the mutated area plus 10 base pairs in each direction. Also, the length of the amplicon was required to match the expected size. The area was then compared to the template sequence using a custom script (`GenerateMut`file.py) to compress the sequences to a list of mutations. Identical sets of mutations were then counted (`SeqCount.py`), resulting in a file with the mutations of the sequences and how many times they were read. For each sequencing pool, a cut-off was chosen to eliminate noise from the dataset based on the mutation rate in the 2×10 base pair region immediately down and upstream of the mutated area.

### GMMA

The genotype counts were aggregated per amino acid variant and 226 complementing variants that were not observed in the input library were discarded. No pseudo-counts were used. Variants with any counts above the cut-off values were considered complementing in the binary readout. This resulted in 838 unique amino acid substitutions combined in 14.887 protein variants holding on average of 9.0 substitutions and 18.5% variants that complement growth in CPOP.

Using the mean-field approach, initial stability estimates could be obtained for all 838 unique amino acid substitutions based on the 10,235 variants that did not contain non-sense mutations (4622) or substitutions that like-wise appeared irreversible fatal (30). Of these, relatively few (194) substitutions were observed in only active or only inactive variants.

For the global analysis, a network analysis found that all substitutions were connected, i.e. that no subset of substitutions only occurred together and never with the rest of the substitutions. This test was important because only a connected network of substitutions can inform the global analysis. Only 54 of the 838 unique substitutions only occurred in a single variant, i.e. are hanging substitutions that do not inform the global analysis. The reference stability was estimated to -27.9 kJ/mol. This is substantially less stable than the value of 48 kJ/mol obtained from chemical unfolding which indicates that the absolute scales of stabilities are not directly comparable. Either because the global model is inaccurate in this respect, or because complementation failure in CPOP only requires partial unfolding and not necessarily the complete global unfolding or simply, that the Trx domain is less stable in the fusion. Furthermore, with a binary phenotype we may not expect the absolute stability scale to be accurate and the present study only relies on the ranking of the substitutions (supplementary figure 3), although the absolute scale does seem relevant (figure 2).

The GMMA error analysis was slightly different from previously described^23^. We did not obtain errors on the estimated stability effects since uncertainties were not determined in the binary readout from the CPOP screen. For filtering of inaccurately estimated effects, we simply required that a substitution should be observed in at least 40 different variants and have a fitting standard uncertainty of 6.3 kJ/mol or better. This resulted in 293 accurately estimated effects.

Requiring that a substitution has been observed in at least 40 variants is rather conservative compared to previous applications of GMMA and was here selected for robustness against the potentially noisy experimental data. Additionally, 81 substitution effects estimated to destabilize more than the reference stability but with higher uncertainty were included as destabilizing resulting in 31 stabilizing, 79 neutral, 264 destabilizing and 462 unknown substitution effects.

### Protein purification

Genes encoding single substitution variants of edF106, derived from the GMMA, were custom synthesized by Twist Bioscience cloned into pET-29b(+) using restriction sites Ndel and XhoI flanking the His_6_ sequence. This resulted in a C-terminal insertion of leucine and glycine before the His_6_-tag. Plasmids were solubilized in TE buffer to 10 ng/µL and 2 µL were used to transform chemically competent BL21 (DE3) cells which were subsequently plated on LB medium with 50 ng/mL kanamycin. Starter cultures were prepared by using single colonies to inoculate 800 µL LB medium with 50 ng/mL kanamycin in a 48 well plate format and incubating over night at 37°C. 2 mL of TB-5052 auto induction medium were inoculated with 20 µL overnight culture and grown for 24 hours at 25°C in 24 deep well plates. TB-5052 is a phosphate buffered medium containing salts and metals for optimized protein expression and a mix of glucose, lactose, and glycerol for auto-induction of the Lac-promoter once the culture as reached appropriate density^39^. Next day, cells were harvested in the plate by centrifugation at 4250 g for 20 minutes. The supernatants were removed and 500 µL lysis buffer pH = 7.0 (50 mM phosphate, 300 mM NaCl, 20 mM imidazole, and 1x BugBuster (10x solution from Novagen) were added to the cell pellets. The plate was incubated while shaking for 25 minutes for lysis before pelleting the insoluble part of the lysate by centrifugation at 4250 g for 40 minutes. The supernatants were transferred to a 96 well filter plate with 250 µL 50% slurry of nickel-NTA beads in each well. After wash with a total of 2 mL buffer with 20 mM imidazole the proteins were eluted in 200 µL buffer with 400 mM imidazole. To remove the imidazole, the IMAC eluates were buffer exchanged on Nap5 columns and the peak fractions were eluted in 200 µL analysis buffer (50 mM phosphate and 150 mM NaCl). 1 µL protein solution was mixed with 19 µL 0.1% TFA for mass spectrum analysis to confirm mutant identity.

### Protein stability measurements

Two-dimensional denaturation and renaturation of the proteins was measured using the Prometheus NT.48 (NanoTemper) using a heating and cooling ramp of 1°C/min. For the preliminary estimation of stabilities, twelve 40 µL samples with equally spaced guanidine hydrochloride (GuHCl) concentration were prepared from two 250 µL solutions, one having 6 M GuHCl, with the same protein concentration (>=5 µM) using a pipetting robot (1000G Andrew Alliance) for consistency. After loading the samples into the capillaries, the ends were sealed with high vacuum grease (Dow Corning) to avoid evaporation during the experiment. Folding and refolding curves were acquired. 5 µL sample without GuHCl were analysed on SDS-PAGE to check that the proteins were pure. The data obtained were fitted using 211103_dTrx_stability.ipynb based on the `ProteinUnfolding2D.py` python module^29^. A single scalar parameter was included to account for the loss of intensity in the refolding. After fitting individual m-values to each dataset in the initial fit, the average m-value was subsequently used for all dataset to get more reliable ΔΔG-estimates. Samples without GuHCl were not used for the fits for all datasets, and for MM3, MM6, and MM9 all samples with more than 2 M GuHCl were excluded from the fits. For absolute estimation of the stability of MM9 and eMM9, samples were prepared with increased density in the transition region.

### Crystallization of MM9 and eMM9

The MM9 construct was concentrated to 10 mg/ml for crystallization experiments using the Hampton screen I and II (Hampton Research). Crystal drops were mixed using 1 uL of protein and 1 uL precipitant solution in 24-well plate as hanging drops on siliconized glass cover-slides. The wells were sealed with vacuum grease (Dow Corning high-vacuum silicone). Plates were incubated at room temperature. Initial crystals of MM9 appeared after approximately a month and grew to a maximal size of 100×100×300 µm, crystal condition: 0.2M NaOAc, 0.1M Tris-HCl (pH 8.5) and 30% polyethylene glycol 4000 (PEG4000). Crystals were harvested using mounted CryoLoops (Hampton Research) and flash frozen in liquid nitrogen. Cryo protection was performed by quick dipping the crystal 0.1 M NaOAc, 0.05 M Tris-HCl (pH 8.5), 15 % polyethylene glycol 4000 (PEG4000) and 20% Glycerol. The data were collected from crystals cooled to 100 K on a PILATUS detector at BioMax (MAX-IV, Lund, Sweden). A full sweep of 360° data was collected with an oscillation degree of 0.1°, with 0.050s exposure, at 12650 eV. Complete data set was processed from 200° (2000 images) with xia2^40^ using the dials pipeline option to account for the weak ice rings (see Supplementary Table 4).

The eMM9 crystals were obtained in a similar procedure, but initial crystals of eMM9 appeared in seven days and grew to a maximal size of 100×100×300 µm. The best eMM9 crystals were grown using a reservoir solution of 0.2 M ammonium sulfate, 0.1 M sodium acetate, pH 4.6, 25% w/v Polyethylene glycol 4.000. Crystals were harvested using mounted CryoLoops (Hampton Research) and flash frozen in liquid nitrogen, cryo protection was performed by quick dipping the crystal in 0.2 M ammonium sulfate, 0.1 M sodium acetate, pH 4.6, 25% w/v Polyethylene glycol 4.000, 20% Glycerol. The data were collected from crystals cooled to 100 K on a PILATUS detector at BioMax (MAX-IV, Lund, Sweden). A full sweep of 360° data was collected with a oscillation degree of 0.1°, with 0.050s exposure, at 12650 eV. Complete data set was processed from 180° (1800 images) with xia2 using the dials pipeline option to account for the weak ice rings (see Supplementary Table 4).

Molecular replacement using the program Phaser^41^ was used to solve the phases using the structure of dF106 (PDB: 5j7d) as an initial search model. The initial model was build using the AutoBuild wizard within the PHENIX package^42^, and for eMM9 the twinning operator h,-h-k,-l was used to account for the crystal twinning. The structure was further manually refined using phenix.refine^43^. Final model building was performed in Coot^44^. Data collection and refinement statistics are summarised in Supplementary Table 4.

### Calculation of Rosetta stabilities

These calculations were carried out as described previously^45^. Briefly, we used the Cartesian ΔΔG protocol^31^ and the X-ray structure of dF106 (PDB: 5J7D). During the initial relaxation of the structure, a res-file was used to introduce L11P and D83V in order to obtain a model of edF106 to be used with the cartesian_ddg application.

### Calculation of lbsDCA conservation scores

These calculations were carried out as described previously^45^. Briefly, we used a statistical analysis of a multiple sequence alignment (MSAs) generated by HHBlits^46^ using the sequence edF106 as target. We used a modified version of the lbsDCA^47^ that includes both positional and pairwise conservation of amino acids. While originally designed to identify contacts between residues, we use the energy potential generated by the algorithm to evaluate the log-likelihood difference between the wild type and the variant sequences.

