## Supplementary material for "Increasing protein stability by inferring substitution effects from high-throughput experiments"

### Supplementary materials

#### Software

Python 3.7.2 was used together with packages: biopython 1.73[48], matplotlib 3.0.3[49], matplotlib-venn 0.11.5, numpy 1.16.2[50], and pandas 0.24.1[51]. ChimeraX was used to create figures of molecular models[52]. Inkscape was used to create figures.

#### Sequences

”Doped” oligonucleotides (Upper case letters symbolize mutagenized positions):

```
> oligo [48:72]
accccuACTGTCGCATTTTTCAAAAATGGCGAGGTCAAGAGCGTTCTGATTGGCGCGATTCC
AAAAGACCAGCTG
> oligo [74:97]
aggggu GGTGGTGATATTGTACTTTTTCGCGATCTCCGGTGCCTTGTTCGTGTCCAGCTCGA
AAACCATGATCAGAC
```

Primer for amplifying the “doped” oligonucleotides alone:

```
>[48:72]
ACCCCUACTGTCGCATTTTTCAAAAATGGCGAGGTCAAGAG
>[74:97]
AGGGGUGGTGGTGATATTGTACTTTTTCGCGATCTCC
```

1st PCR Illumina amplicon primers (Upper case letters is the region representing Nextera adapters and lower case is the region complementary with the plasmid)

```
>Forward
TCGTCGGCAGCGTCAGATGTGTATAAGAGACAGccgtaaagataacgacgaagaggc
>Reverse
GTCTCGTGGGCTCGGAGATGTGTATAAGAGACAGtcgatgaactgacgttggtacgg
```

edF106 gene:

```
ATGGTACTGGATGTAACGAAAGATCACTGGCTGCCCTACGTATTACTCGCTCAACTGCCGGT
CATGGTGTTGTTCCGTAAAGATAACGACGAAGAGGCCAAGAAGGTTGAGTATATTGTGCGCG
AACTGGCGCAGGAATTTGACGGTCTGATCATGGTTTTCGAGCTGGACACGAACAAGGCACCG
GAGATCGCGAAAAAGTACAATATCACCACCACCCCGACTGTCGCATTTTTCAAAAATGGCGA
GGTCAAGAGCGTTCTGATTGGCGCGATTCCAAAAGACCAGCTGCGTGATGAAATCCTGAAAT
ATCTGGGTCACCATCATCACCATCAC
```

#### Supplementary tables

| Sample | Total reads | Reads passed (%) | Unique DNA sequences after cut-off | Unique protein sequences after cut-off |
|---|---|---|---|---|
| [48:97] input | 1,357,174 | 356,577 (26.3) | 15018 | 14978 |
| [48:97] 30deg | 1.494,800 | 654,314 (43.8) | 3009 | 2980 |

Supplementary Table 1: Sequencing data from the libraries and for the filtering of sequences based on how many times they were observed. The lower fraction of reads passing the filters for the input libraries in part reflect species that were cloned incorrectly and therefore did not have the correct length.

| Sample | [48:72] area | [74:97] area | Total | Not mutated |
|---|---|---|---|---|
| [48:97] input | 6.61 | 6.74 | 13.40 | 0.05 |
| [48:97] 30deg | 5.34 | 4.90 | 10.27 | 0.03 |

Supplementary Table 2: The average mutations per sequence in each of the samples. The mutation G148T (at the DNA level) is omitted from this count because it was a technicality of the cloning procedure. Mutations in the regions originating from the mutated oligonucleotides are counted separately. The [74:97] oligonucleotide is slightly less mutated than then [48:72]. The last column shows that very few mutations were seen outside of the mutated area.

| Protein | ΔG (25°C) | m-value | Tm | ΔHs | ΔCp | $X^2$ | ΔΔG |
|---|---|---|---|---|---|---|---|
| edF106 | 46.83 | 12.67 | 109.25 | 477.63 | 5.81 | 2.29E+08 | 0.00 |
| M51K | 49.43 | 12.67 | 103.54 | 555.29 | 7.52 | 1.68E+08 | -2.61 |
| M51R | 47.32 | 12.67 | 105.39 | 507.02 | 6.55 | 2.40E+08 | -0.49 |
| M51T | 42.39 | 12.67 | 102.24 | 464.82 | 6.22 | 4.41E+08 | 4.44 |
| E54V | 53.65 | 12.67 | 113.80 | 529.29 | 6.12 | 6.63E+08 | -6.82 |
| E54Y | 45.26 | 12.67 | 110.98 | 448.62 | 5.29 | 2.00E+07 | 1.57 |
| L55V | 51.01 | 12.67 | 110.65 | 502.96 | 5.91 | 1.18E+08 | -4.18 |
| T57I | 52.12 | 12.67 | 119.04 | 485.83 | 5.23 | 1.52E+08 | -5.29 |
| K79H | 44.29 | 12.67 | 115.03 | 432.14 | 4.93 | 3.37E+07 | 2.54 |
| V83L | 56.21 | 12.67 | 108.30 | 579.21 | 7.14 | 2.96E+08 | -9.38 |
| L87F | 51.68 | 12.67 | 110.77 | 504.76 | 5.88 | 5.91E+07 | -4.85 |
| I88S | 49.38 | 12.67 | 103.60 | 527.14 | 6.86 | 2.98E+08 | -2.55 |
| I88T | 49.72 | 12.67 | 109.64 | 488.73 | 5.76 | 2.03E+08 | -2.90 |
| P92R | 49.94 | 12.67 | 110.75 | 499.32 | 5.93 | 9.08E+08 | -3.11 |
| P92S | 55.28 | 12.67 | 110.16 | 549.43 | 6.52 | 4.06E+08 | -8.45 |
| P92T | 53.98 | 12.67 | 117.50 | 499.43 | 5.39 | 5.45E+08 | -7.15 |
| MM3 | 56.12 | 12.67 | 113.39 | 532.98 | 5.99 | 9.19E+07 | -9.29 |
| MM6 | 60.95 | 12.67 | 132.59 | 487.43 | 4.34 | 6.97E+07 | -14.13 |
| MM9 | 67.58 | 12.67 | 143.97 | 499.94 | 3.98 | 5.33E+07 | -20.75 |
| MM9_0 | 67.68 | 12.67 | 170.86 | 431.53 | 2.73 | 1.49E+09 | -20.85 |
| MM9_1 | 69.76 | 12.67 | 151.34 | 505.07 | 3.84 | 1.30E+09 | -22.93 |
| MM9_2 | 69.81 | 12.67 | 145.86 | 527.09 | 4.24 | 1.49E+09 | -22.98 |
| MM9_3 | 68.38 | 12.67 | 149.36 | 511.37 | 4.02 | 6.32E+08 | -21.55 |
| eMM9_0 | 67.97 | 12.67 | 157.09 | 470.13 | 3.36 | 2.21E+08 | -21.14 |
| eMM9_1 | 70.34 | 12.67 | 145.17 | 530.95 | 4.28 | 1.53E+09 | -23.51 |
| eMM9_2 | 70.24 | 12.67 | 146.79 | 519.73 | 4.09 | 7.43E+08 | -23.42 |
| eMM9_3 | 69.27 | 12.67 | 148.92 | 510.02 | 3.97 | 4.27E+08 | -22.44 |

Supplementary Table 3: Fitted parameters of combined temperature and denaturant unfolding of purified proteins for Figure 2a, b and c. The m-value was kept constant (see Methods). ΔG (25°C) and ΔΔG values were derived from the other fitted parameters. ΔG, ΔH and ΔΔG are in kJ/mol, the m-values are in kJ/(mol·M), $T_m$ is in K, and $\Delta C_p$ in kJ/(mol·K).

Supplementary Table 4: Crystal structure statistics

| | MM9 (PDB: 7Q3J) | eMM9 (PDB: 7Q3K) |
|---|---|---|
| Wavelength | 0.98 | 0.98 |
| Resolution range | 25.93 - 1.9 (1.968 - 1.9) | 47.82 - 2.0 (2.072 - 2.0) |
| Space group | C 1 2 1 | P 31 |
| Unit cell | 58.89 45.66 72.88 90 92.16 90 | 71.52 71.52 75.22 90 90 120 |
| Total reflections | 53102 (5297) | 152832 (15197) |
| Unique reflections | 14904 (1490) | 29073 (2918) |
| Multiplicity | 3.6 (3.6) | 5.3 (5.2) |
| Completeness (%) | 96.53 (95.69) | 99.82 (99.79) |
| Mean I/sigma(I) | 22.28 (2.24) | 8.23 (1.19) |
| Wilson B-factor | 28.66 | |
| R-merge | 0.063 (0.32) | 0.079 (0.87) |
| R-meas | 0.074 (0.37) | 0.0873(0.96) |
| R-pim | 0.039 (0.19) | 0.038 (0.42) |
| CC1/2 | 0.997 (0.755) | 0.998 (0.916) |
| CC* | 0.999 (0.928) | 1 (0.978) |
| Reflections used in refinement | 14893 (1487) | 29035 (2917) |
| Reflections used for R-free | 759 (74) | 1425 (150) |
| R-work | 0.19 (0.28) | 0.28 (0.39) |
| R-free | 0.24 (0.30) | 0.30 (0.46) |
| CC(work) | 0.96 (0.75) | 0.89 (0.72) |
| CC(free) | 0.91 (0.84) | 0.88 (0.89) |
| Number of non-hydrogen atoms | 1532 | 2513 |
| macromolecules | 1445 | 2455 |
| ligands | 28 | 15 |
| solvent | 75 | 43 |
| Protein residues | 176 | 302 |
| RMS(bonds) | 0.012 | 0.004 |
| RMS(angles) | 1.17 | 0.91 |
| Ramachandran favored (%) | 98.84 | 94.86 |
| Ramachandran allowed (%) | 1.16 | 4.11 |
| Ramachandran outliers (%) | 0.00 | 1.03 |
| Rotamer outliers (%) | 0.00 | 2.94 |
| Clashscore | 5.09 | 17.10 |
| Average B-factor | 39.22 | 71.87 |
| macromolecules | 39.06 | 72.16 |
| ligands | 50.08 | 87.18 |
| solvent | 40.64 | 49.97 |
| Number of TLS groups | 12 | 11 |

Statistics for the highest-resolution shell are shown in parentheses.

#### Supplementary figures

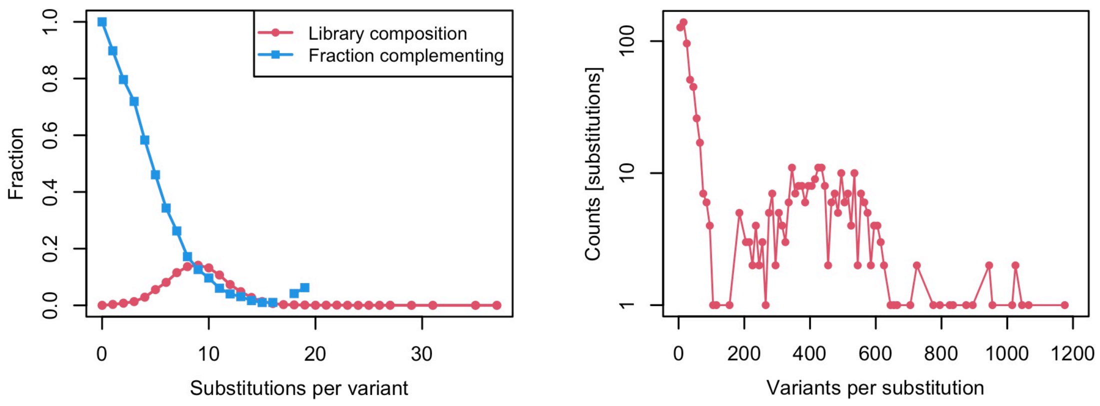


**Supplementary Figure 1: Characterization of the variant library.** Left: The broad distribution of muti-mutants (red) covers the complementation profile (blue) sufficiently to give a total of 19% complementing variants. Variants with 9 substitutions are most abundant and approximately half of the 5-mutants complement in the assay. Right: The library stategy using long mutated primers is shown to result in a relatively narrow peak between 200 and 600 variants per substitution which is a relatively homogeneous amount of data per estimated parameter in GMMA. Effects of substitutions oberserved in less than 40 different variants are assumed to inaccurately estimated in the error analysis.

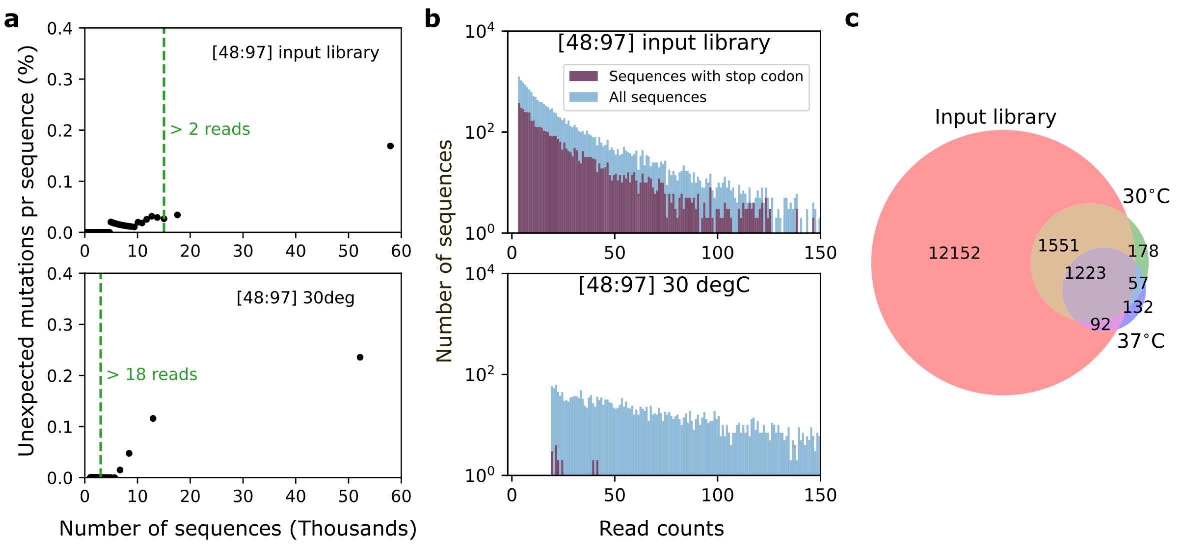


**Supplementary Figure 2: Sequencing data cut-off is choosen to eliminate noise. a,** Plots showing the rationale for choosing cut-offs for the number of sequencing reads required to remove sequencing noise. The metric used to determine the minimum observations of a sequence required to be included in the downstream analysis (cut-off level) is average number of mutations in the 10 base pairs immediately up- and downstream of the region intentionally mutated. Here, these are termed "unexpected mutations". The rate of such mutations were recorded for each cut-off value (>1,>2,>3…>N reads) plotted as single points from right to left (black dots) on the graphs, based on the remaining number of sequences (x-axis). Green text and vertical bars indicate the cut-offs chosen for the analysis. **b,** Sequences with stop codons were highly depleted after selection. Histograms of readcounts for each sequence identified by MPS from the library [48:97] in blue, with the fraction of sequences containing a stop codon in red. **c,** Venn diagram showing the amount of reads found in the different libraries before, and after selection at 30°C and 37°C.

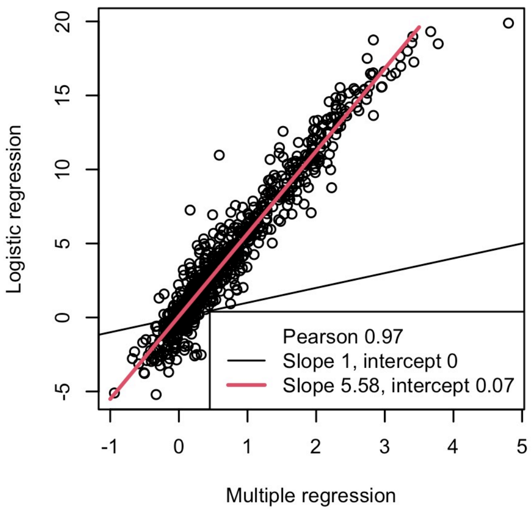


**Supplementary Figure 3. Using binary data for GMMA is appropriate.** GMMA of GFP variants from a previous study (Johansson et al., 2021). The plot shows the correlation between estimated stability effects for the original continuous fluorescence readout (multiple regression) and a binary readout generated from the same data (logistic regression). The correlation shows that for GFP, the ranking and zero-point (intercept) is preserved. The results also show that the scale of GMMA changes when the assay is made binary.

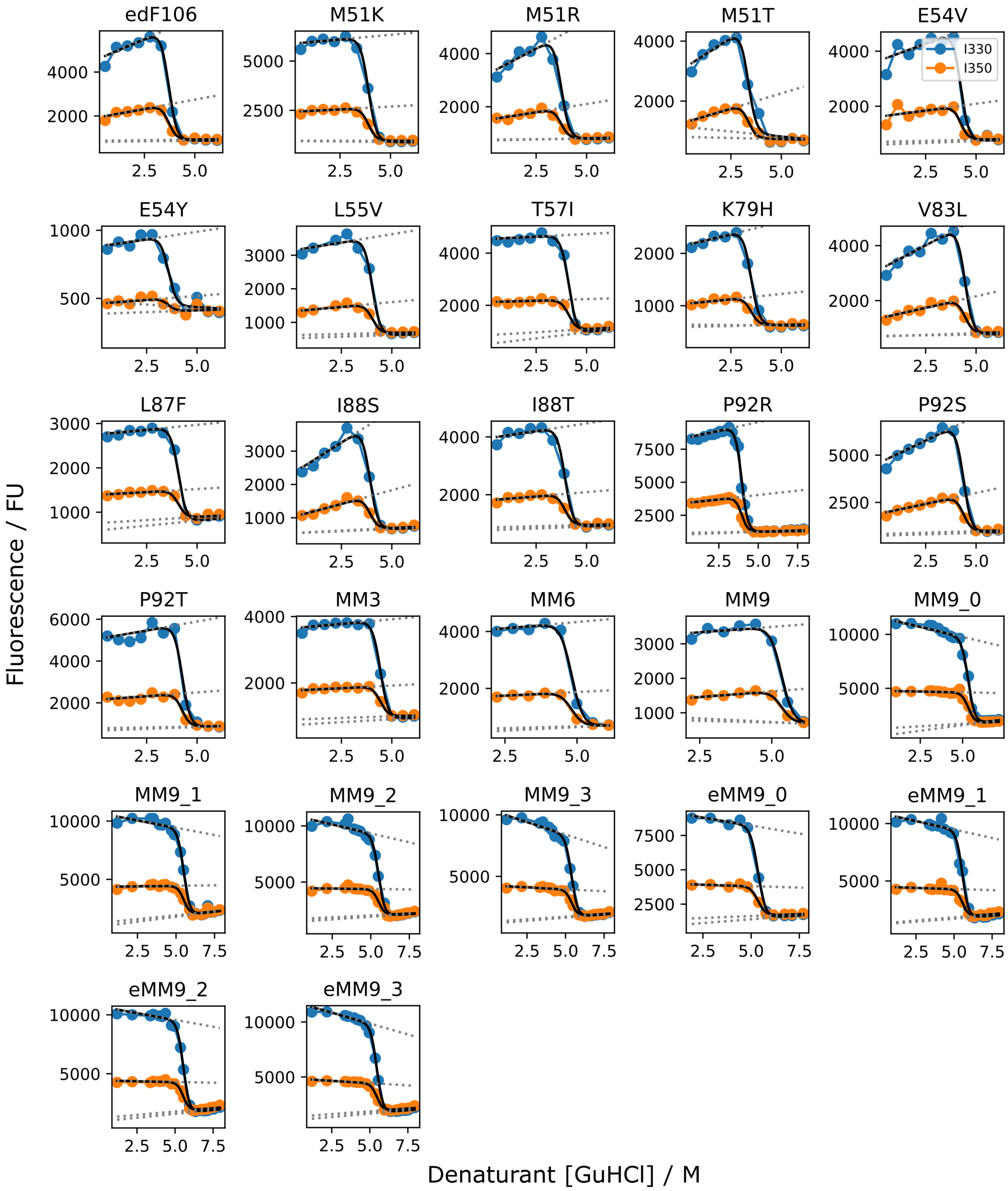


**Supplementary Figure 4: Isotherms from two-dimensional fitting of variants.** Raw data for fitting global denaturation using denaturant and temperature shown only in the denaturant dimension. The data from the fluorescence intensity at 330 and 350 nm obtained on the Prometheus NT.48 for folding and refolding was fitted globally. The baselines of the fits are shown in grey dots

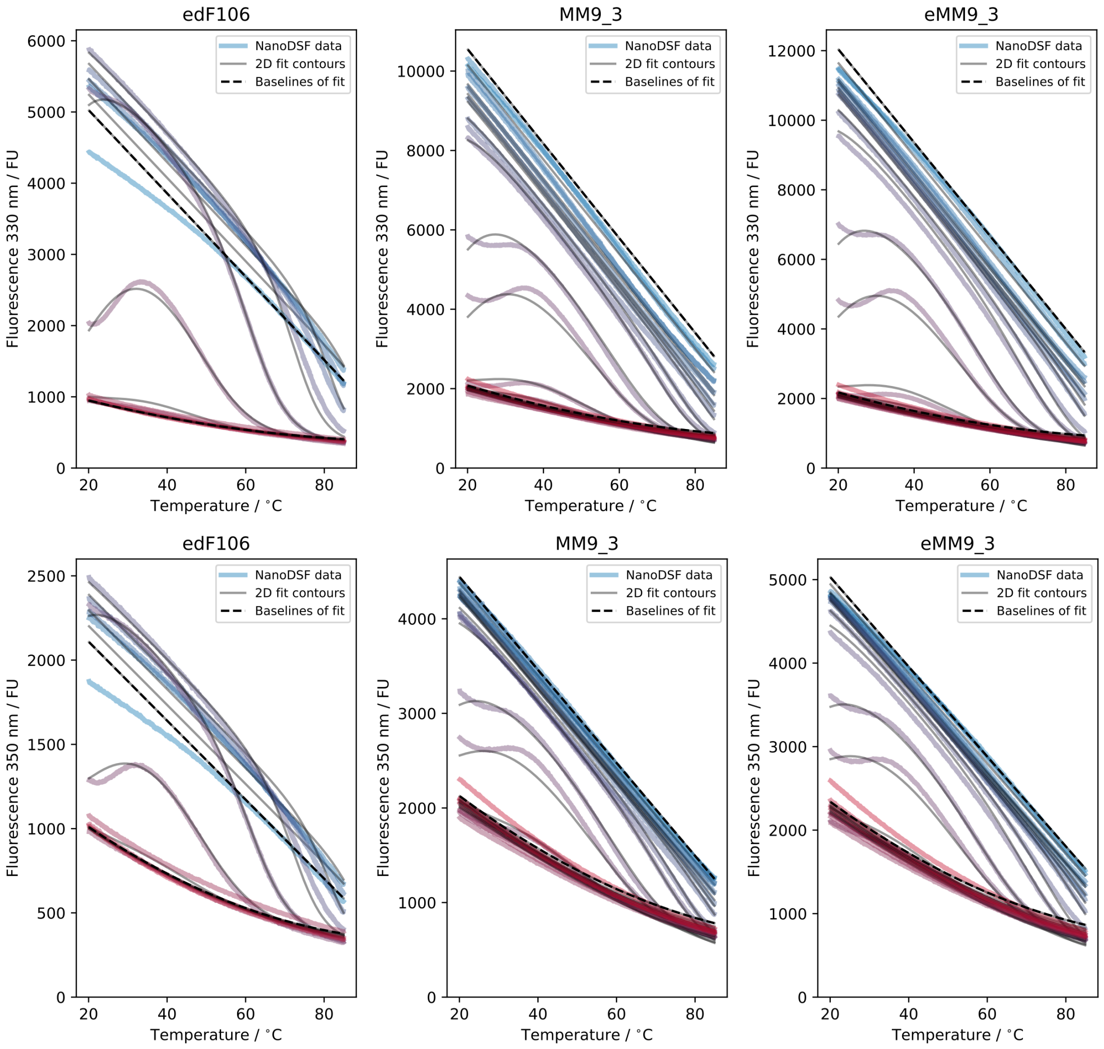


**Supplementary Figure 5: Fit evaluation for two-dimensional unfolding.** Full two-dimensional fits of edF106, MM9, and eMM9 are shown in the temperature dimension. The fluorescence intensities during unfolding at 330 nm and 350 nm when excited at 280 nm in NanoDSF are fitted. The samples are coloured from blue to red with increasing concentration of GuHCl: 0.6-6.1 M for edF106, 1.1-7.1 M for MM9, and 1.9-7.7 M for eMM9. Fitting was done globally on both traces of folding and the corresponding contours are shown in black with the highest and lowest concentration of GdnHCl shown as interrupted lines.

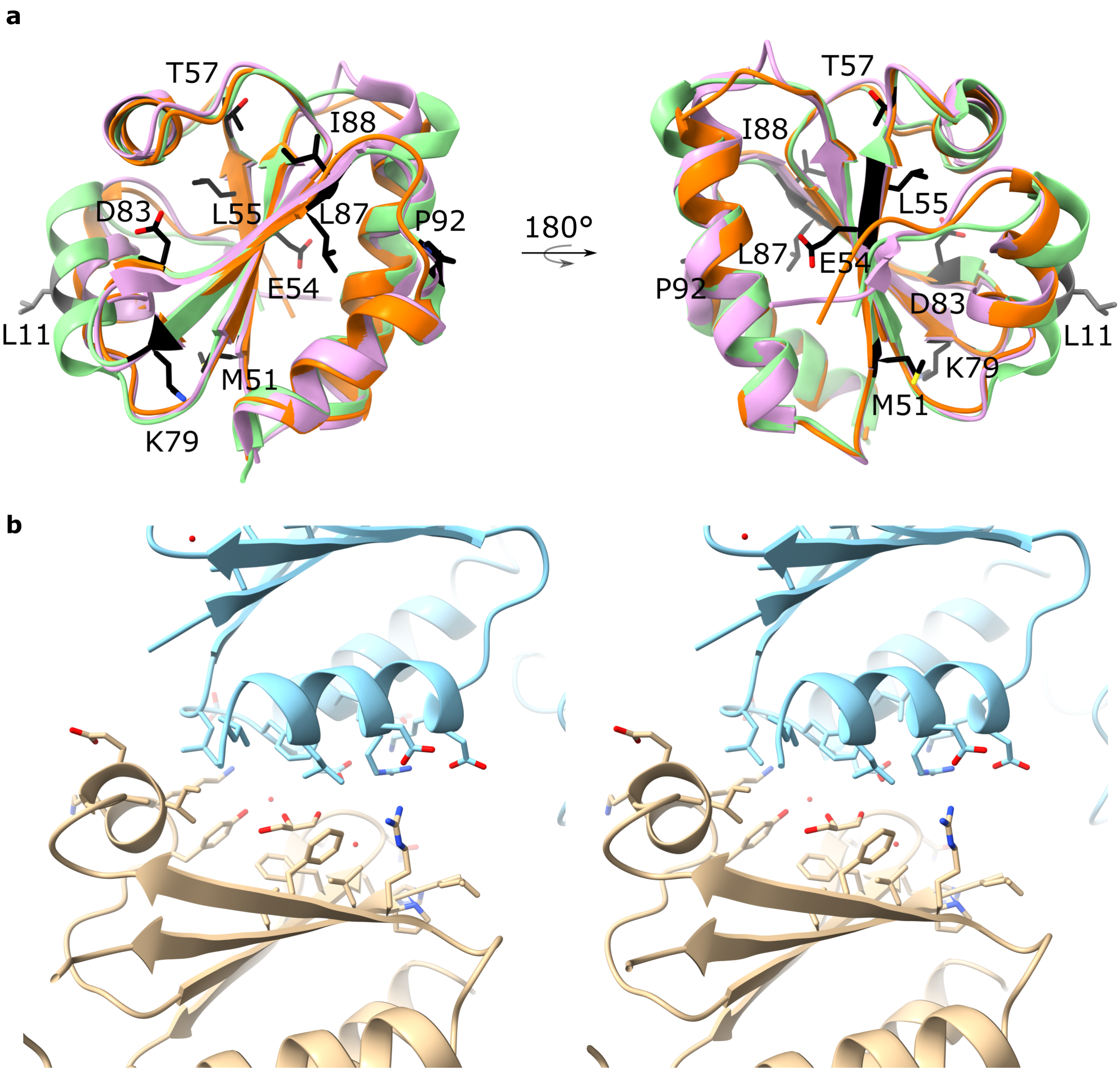


**Supplementary Figure 6: Crystal structures.** a, Mutated residues to MM9 are drawn in stick representation and coloured in black. The residue L11 is coloured in grey because it was one of the original mutations in dF106 to make edF106. dF106 (PDB: 5J7D) in green, eMM9 (PDB: 7Q3K) in orange and the original spinach Trx design template (PDB: 1FB0) in purple. b, Stereo view of the crystal contact formed in the MM9 structure (PDB: 7Q3J) in the pocket normally occupied by the N-terminal. Here, the N-terminal has displaced and instead α-helix 4 and parts of α-helix 2 forms a symmetry contanct involving a solvent glycerol molecule.
